# Loss of LRP1 in adult neural stem cells impairs migration to ischemic lesions

**DOI:** 10.1101/2022.08.17.504194

**Authors:** Kristi Dietert, Swetha Mahesula, Sheetal Hegde, John Verschelde, Pamela Reed, Shane Sprague, Erzsebet Kokovay, Naomi L. Sayre

## Abstract

After ischemia, cells in the brain parenchyma upregulate stromal derived factor 1 (SDF1), driving chemokine receptor CXCR4-mediated migration of adult neural stem cells from the subventricular zone (SVZ) to the ischemic injury. We discovered a novel regulator of CXCR4 in neural stem cells, low-density lipoprotein receptor related protein 1 (LRP1). We employed a tamoxifen-inducible Nestin-Cre to drive expression of a tdTomato reporter and also knockout floxed LRP1 in adult mice and then subjected mice to middle-cerebral artery occlusion. Examination 2 weeks post-stroke reveals a loss of tdTomato positive cells localizing from the SVZ to the lesion. We show that loss of LRP1 disrupts CXCR4-mediated neural stem cell migration *in vitro*, which is likely driven by LRP1-mediated loss of CXCR4 expression *in vivo*. Altogether, our results suggest that LRP1 is a novel regulator of CXCR4 in neural stem cells.

**Highlights:**

- LRP1 KO in adult neural stem cells disrupts migration to ischemic lesions *in vivo*.
- LRP1 KO in adult neural stem cells disrupts migration towards SDF1 *in vitro*.
- LRP1 positively regulates expression of CXCR4 in adult neural stem cells.

**eTOC blurb:** Adult neural stem cells can home to ischemic brain injury and are considered an important part of the repair process after stroke. However, little is known about what molecules help drive this response. The authors discovered that LRP1 is a novel regulator of CXCR4, which is essential for neural stem cell migration to ischemic injury.

## Introduction

Stroke is a leading cause of death and a primary cause of disability worldwide^1,2^. An estimated 87% of strokes are ischemic strokes, which arise from occlusion of blood flow^3^. Currently, subacute therapeutics capable of mitigating post-stroke damage are lacking, necessitating improved understanding of mechanisms which drive post-stroke damage and repair.

Multiple signals are released in the ischemic lesion after stroke; neural stem cells (NSCs) resident within the subventricular zone (SVZ) niche are prime targets for such signals. After stroke, NSCs become activated and proliferate to expand the progenitor pool in the SVZ^9,4,5^. The chemokine SDF1 (stromal cell-derived factor 1) is secreted by astrocytes and endothelial cells in the ischemic lesion6, causing neuroblasts to divert from the rostral migratory stream (RMS) and instead migrate toward the lesion^9,7–11^. Once there, they give rise to a limited number of neurons^12–14^ and secrete trophic factors^13,15–17^. Ultimately, NSC migration toward the lesion is considered protective^9,11,13–17^, because genetic ablation of neural progenitors increases lesion sizes after stroke^15,18,19^.

The ability of NSCs to proliferate, migrate, and differentiate in a neuroprotective manner depends on dynamic and responsive cellular signaling. The low-density lipoprotein receptor related protein 1 (LRP1) is a key, albeit underappreciated, player in signal regulation. LRP1 is implicated in the binding/trafficking of over 40 proteins in multiple cell types^20^. When bound to ligand/plasma membrane proteins, LRP1 undergoes receptor mediated endocytosis to traffic bound protein for degradation^21–24^. LRP1 can also be cleaved by γ-secretase, translocating to the nucleus to regulate gene expression^25–28^. Currently, the degree to which LRP1 influences signaling in NSCs is unclear, but many pathways relevant to NSC biology are regulated by LRP1^29–31^.

We sought to test a role for LRP1 in regulation of NSCs in the post-stroke milieu. We discovered that knockout of LRP1 in adult NSCs impaired migration toward ischemic lesions concomitant to a loss in CXCR4 expression. Altogether, our results suggest a heretofore undiscovered player in the regulation of CXCR4 in NSCs, which could have broad implications toward NSC physiology in health and disease.

## Results

### Loss of LRP1 in adult NSCs impairs localization to ischemic lesions

We employed middle cerebral artery occlusion (MCAO) to create a transient (30-minute), hemispheric ischemic stroke via 75% reduction in cortical blood flow. Blood flow returned to normal levels after removing the occlusion (**Fig. 1A**). We subjected Control (Nestin-^CreERt2^:tdTomato:LRP1^+/+^) and LRP1-KO (Nestin-Cre^ERt2^:tdTomato:LRP1^fl/fl^) mice to MCAO at 3 months of age, 1 month after tamoxifen induction (**Figure 1B**). The modified neurological severity score (mNSS) was used to test for functional impairment at 6 hours and 3 days post-stroke. Regardless of genotype, mice exhibited similar cortical blood flow reductions (**Figure 1C**) and mNSS score (**Figure 1D**), suggesting that lesions were of similar severity in the acute period post-stroke.

**Figure 1.**
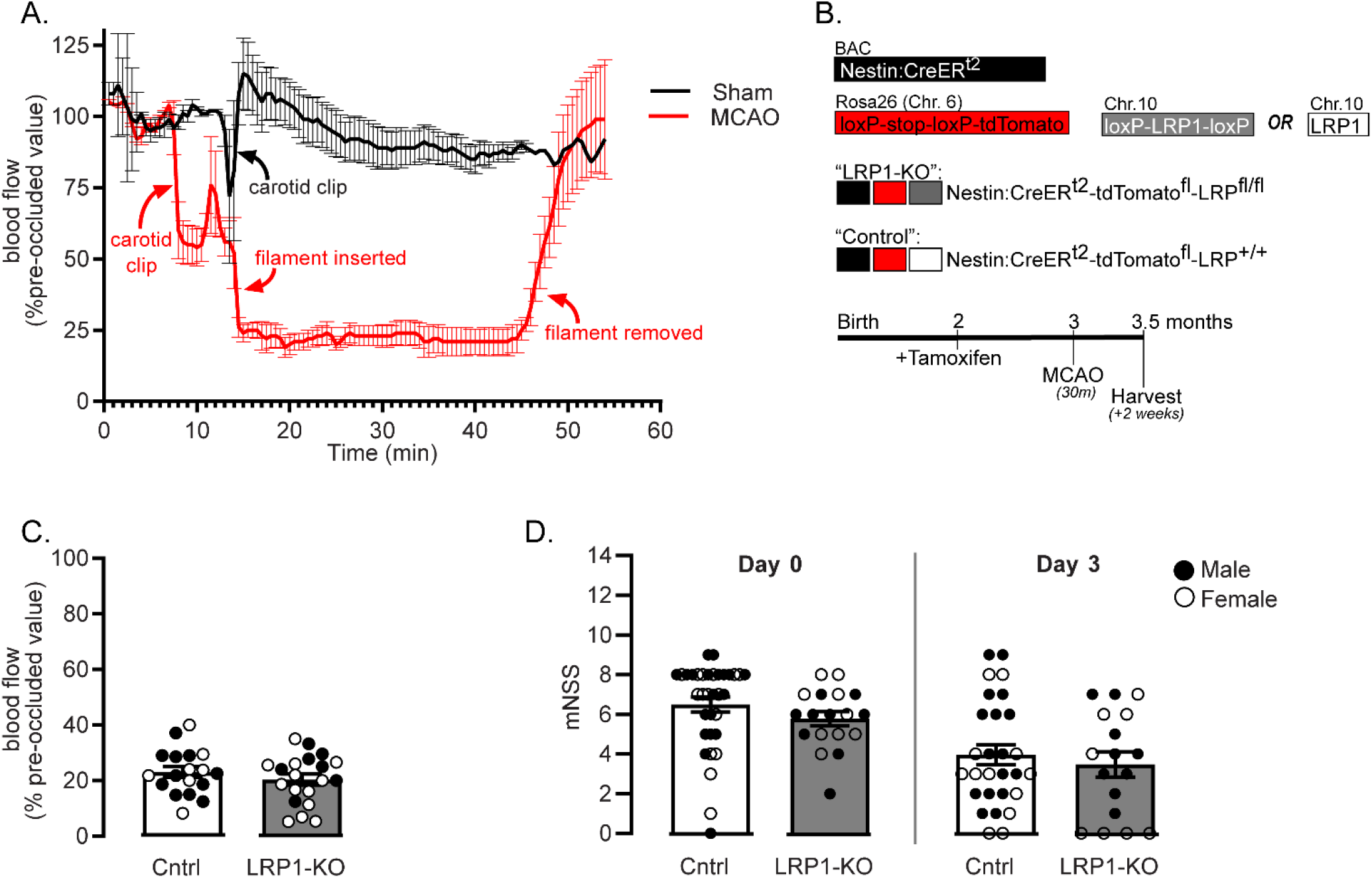
MCAO in inducible NSC-specific LRP1 KO mice. **(A)** Cortical perfusion was visualized via laser-Doppler in sham (black) and occluded (red) mice (n=4/group) **(B)** Nestin:Cre^ERt2^:tdTomato mice were crossed with LRP1^fl/fl^ mice to create Nestin-Cre^ER^:td-tomato:LRP1^fl/fl^ mice (LRP1-KO) or Nestin-Cre^ER^:td-tomato:LRP1^+/+^ (Control). Mice were subjected to MCAO at 3 months of age, 1-month after tamoxifen. **(C)** Cortical blood flow was measured in Control (Cntrl, white) or LRP1-KO (gray boxes) (n=18-20) and expressed as a percentage of pre-occluded blood flow. **(D)** mNSS at 6 hours and 3 days post stroke in n=17-34 mice/group. Results are averages ± SEM. Significant differences were tested using ANOVA followed by Tukey’s HSD.

Mice were harvested 2 weeks post-MCAO, and the distance of tdTomato positive cells that travelled from the SVZ toward the striatal lesion was measured. NSCs lacking LRP1 had reduced localization toward the ischemic lesion compared to controls (**Fig. 2A-E**, p<0.0001). To ascertain if the migration defect was due to a general inability to migrate, we examined if tdTomato positive cells were found in the olfactory bulb of non-stroked mice. Regardless of genotype, tdTomato positive cells could be found along the RMS and in the olfactory bulb (**Fig. 2F,G**).

**Figure 2.**
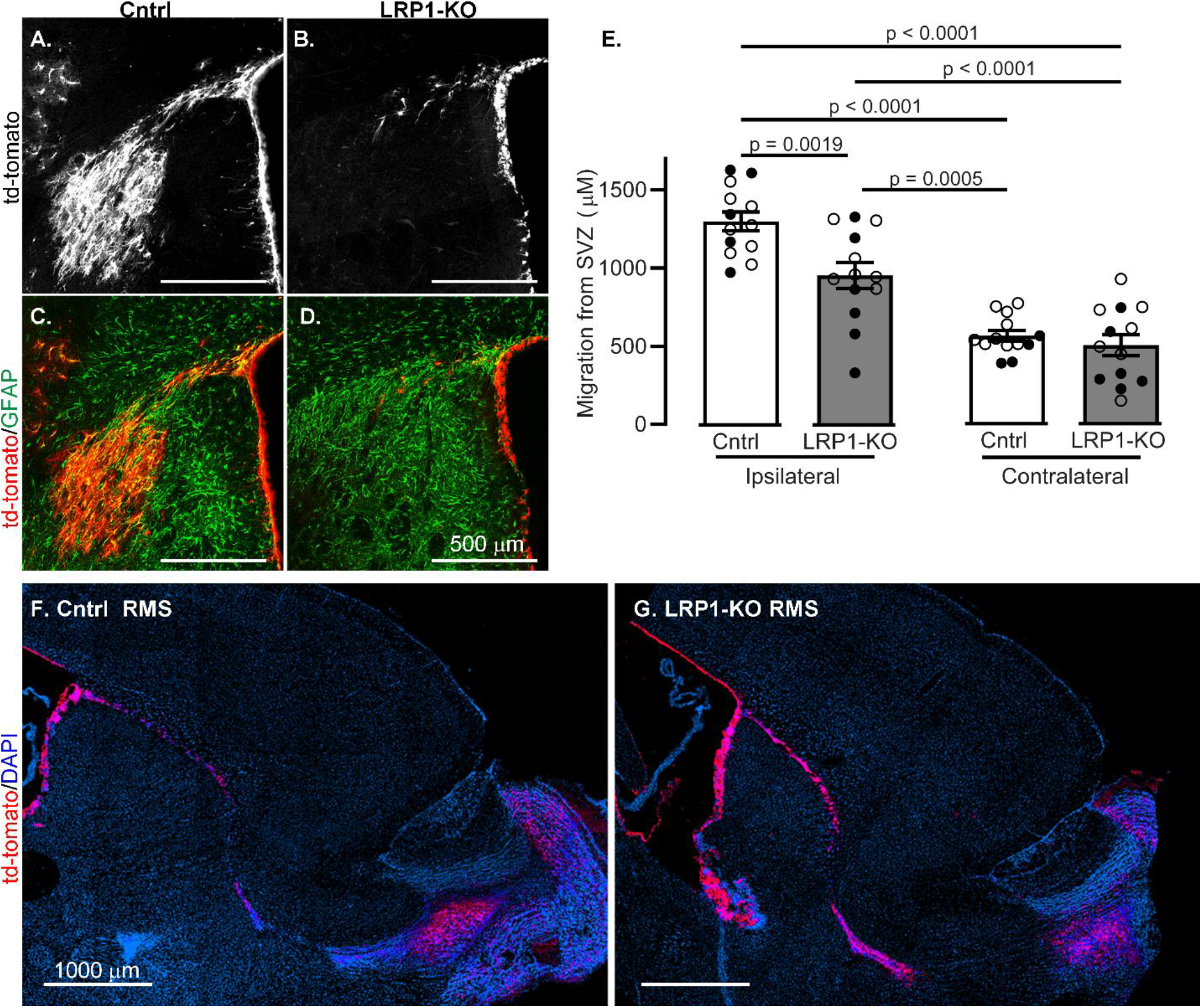
LRP1-KO in adult-born neuroblasts causes migration defects after ischemia. Coronal sections show representative image of tdTomato+ cells (white) 2 weeks after MCAO in **(A)** Control and **(B)** LRP1 KO mice. TdTomato images (red) were merged with GFAP-labelled imaged (Green) to show post-stroke glial scarring from **(C)** Control and **(D)** LRP1-KO mice. **(E)**. Quantification of tdTomato+ cell distance from the SVZ in ipsilateral and contralateral regions (n=13). Results are averages ± SEM. Represented significant differences were tested using ANOVA followed by Tukey’s HSD. Representative images of the RMS in sagittal sections of **(F)** Control and **(G)** LRP1-KO mice.

### LRP1 knock-out causes loss of CXCR4

We tested if CXCR4 expression was affected by LRP1-KO, as this pathway is necessary for neuroblast migration to ischemic lesions^16,19,43^ but is less critical for directed migration along the RMS^32,33^. We tested expression of CXCR4 and found only 30% of CXCR4 labeling in the red tdTomato positive cells in LRP1KO compared to Control mice (p<0.02, **Fig. 3A-I**). We also measured CXCR4 in the NSCs of non-stroked mice and found mRNA expression reduced to approximately 5% of control in LRP1KO cells mice (p=0.013, **Fig. 3J**).

**Figure 3.**
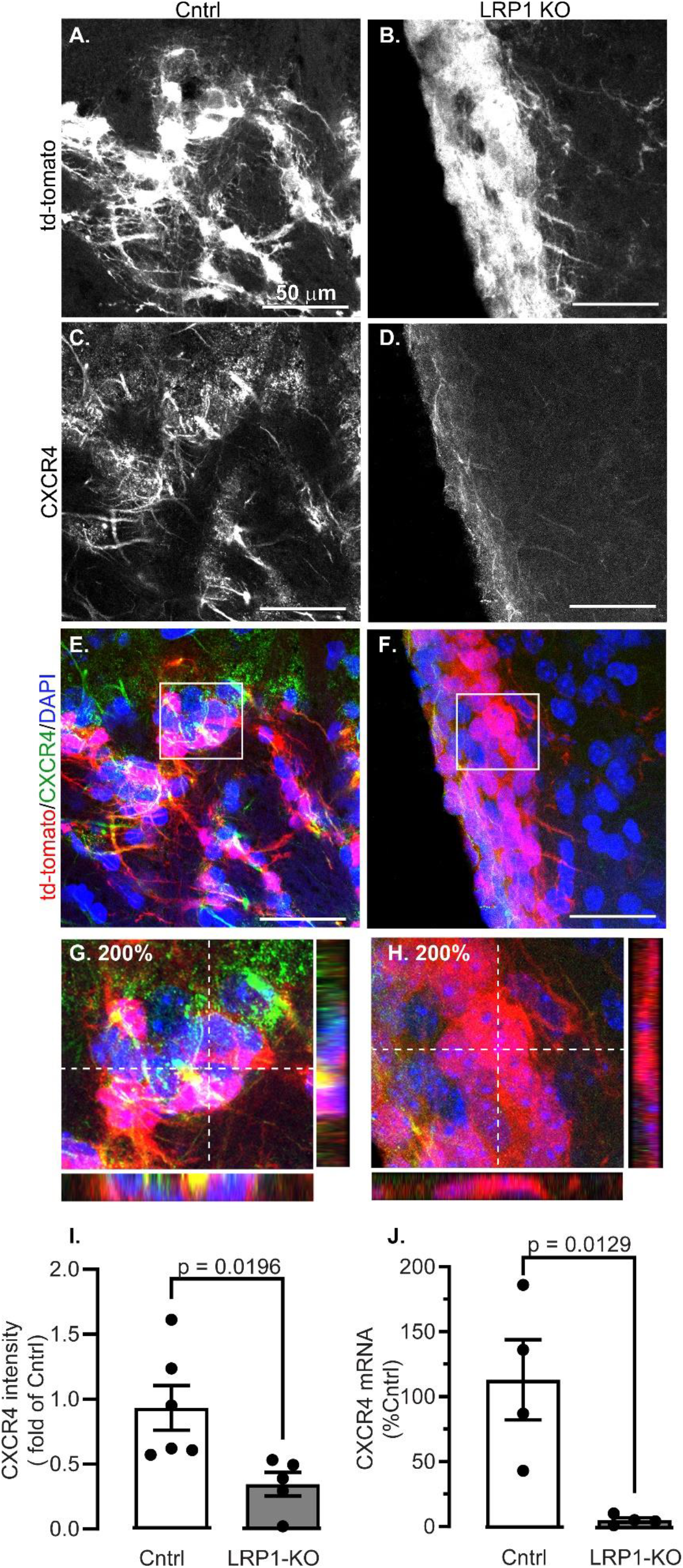
LRP1 KO decreases CXCR4 expression in adult NSCs. Representative images are shown from SVZs in **(A, C, E, G)** Control (Cntrl) and **(B, D, F, H)** LRP1-KO mice 2-weeks after MCAO of **(A,B)** tdTomato+ cells, **(C,D)** CXCR4 immunostained cells, **(E, F)** Channels were merged to show tdTomato (red), CXCR4 (green) and nuclear DAPI (blue). Areas in the white box are magnified in **(G, H)** to show orthogonal views. **(I)** Quantification of CXCR4 intensity in tdTomato+ cells (n=5-6 mice/group). **(J)** qRT-PCR of CXCR4 was in tdTomato+ sorted cells (n=4). Results are averages ± SEM. Significant differences were tested using ANOVA followed by Tukey’s HSD and are represented on the graph.

### LRP1 knock-out impairs migration to SDF1 *in vitro*

The decreased localization of NSCs to the ischemic lesion suggests defective migration toward lesions. To test if LRP1-KO impairs *in vitro* migration, NSCs were cultured from Control and LRP1-KO mice and subject to a transwell migration assay (**Fig 4A**). In the presence of SDF1, significantly greater numbers of tdTomato positive control NSCs migrated through the chamber (p=0.01). In contrast, SDF1 did not increase numbers of migrated tdTomato positive LRP1-KO NSCs (**Fig 4B-F**).

**Figure 4.**
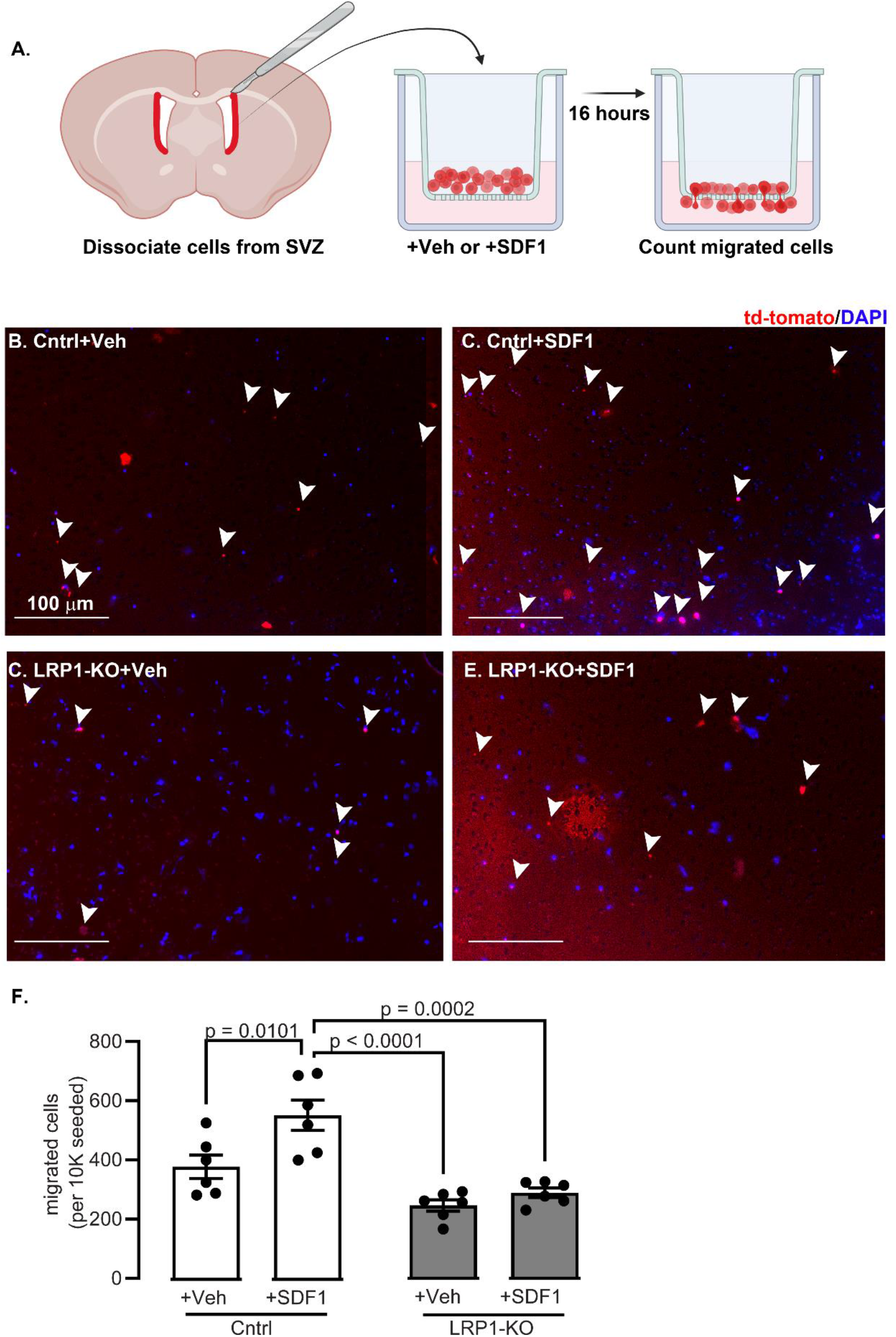
LRP1-KO prevents NSC migratory response to SDF1 *in vitro* **(A)** Schematic of approach for chemotaxis assay with representative images for **(B)** control cells treated with vehicle **(C)** control cells with SDF1 **(D)** LRP1 KO cells with vehicle and **(E)** LRP1 KO cells with SDF1. White arrows show counted migrated cells **(F)** Quantification migrated cells (n=6 wells/group, triplicate independent experiments). Results are averages ± SEM. Significant differences were tested using ANOVA followed by Tukey’s HSD and are represented on the graph.

## Discussion

Currently, little is known regarding a role for LRP1 in NSCs after ischemic damage. We discovered that loss of LRP1 in adult NSCs causes deficits in ischemia-induced migration, which was confirmed *in vitro* and is likely due to a loss of CXCR4.

LRP1 regulates many signaling pathways via interaction with many proteins^20^. To our knowledge, our data is the first to implicate LRP1 in regulating CXCR4. Given the decrease in CXCR4 at both the mRNA and protein levels and a known role for LRP1 as a co-transcriptional activator^25–28^, we hypothesize the regulatory mechanism to be transcriptional. Canonically, NRF-1 is the major positive transcriptional regulator of CXCR4 while Ying Yang 1 is the negative transcriptional regulator, but other transcription factors also alter CXCR4 expression^34,35^. PPARγ downregulates CXCR4 in cancer associated fibroblasts^36^. This is of particular interest because LRP1 can be cleaved by γ-secretase^27^, translocate to the nucleus, and act as co-activator for PPARγ^25^. Thus, it is possible that PPARγ is intermediary in regulation of CXCR4 by LRP1. Future study is warranted.

LRP1 could also regulate CXCR4 through plasma membrane receptor trafficking, which is a known method of CXCR4 regulation^37^, thus LRP1 may modulate this activity. Studies have shown that LRP1 regulates trafficking of CXCR3 in tumor cells^38^, thus it is also a plausible mechanism for CXCR4.

Our observations could have significant implications for tissue recovery after stroke. This report does not include such an analysis, however interrogation into any differences in recovery is underway. Prior reports suggest that blocking SDF1/CXCR4 diminishes stem cell migration to lesions^14,39^ though these studies do not include analysis of the lesions. Other studies suggest that experimental elimination of NSCs is harmful for stroke recovery^15,18,19,40^, thus homing to a lesion could be central to NSCs benefit. Future studies will help test this.

Despite the study limitations, the implications of our discovery in the context of stroke recovery and basic neurobiology are significant. An intricate understanding of mechanisms underlying NSC response during ischemic injury may enable the development of therapeutics that improve natural neurogenic response to mitigate damage and promote recovery. Our discovery could also culminate into advancements of our collective understanding of basic NSC biology. Given that LRP1 knockout is embryonic lethal^41^, it must play a fundamental role in regulation of NSCs, however our understanding is somewhat limited. Safina *et al*. reported the importance of LRP1 in neural stem cells *in vitro--*LRP1 deletion had a negative effect on survival and proliferation and altered differentiation to favor astrocytes and oligodendrocyte progenitor cells^42^. Our studies extend upon such findings and suggest a multi-factorial role of LRP1 on NSC biology *in vivo*. Improved understanding of LRP1 in NSC biology will enhance our ability to acutely harness and regulate neurogenic processes, in turn broadly impacting a number of neurobiological diseases affected by neurogenesis, including schizophrenia^43–45^, mood disorders^44,46,47^, neurodegenerative diseases^48,49^, and epilepsy^50,51^.

## Experimental Procedures

### Animals

All animal procedures were approved by the UT Health Institutional Animal Care and Use Committee (IACUC) in accordance with NIH guidelines. Vivarium conditions included 12h light/dark cycle with *ad libitum* food and water. Adult NSC-specific LRP1 KO mice were generated by crossing: Nestin-Cre^ERT2^ (Jackson stock #016261) to induce Cre recombinase in NSCs, tdTomato Ai14 reporter mice (tdTomato-stop^fl/fl^ stock 007914), and LRP1 floxed mice (stock 012604). KO was induced at 2 months of age with daily IP tamoxifen (100uL of 20mg/ml dissolved in corn oil) administered for 5 days. At 2 weeks post-stroke, mice were euthanized via 5% isoflurane/oxygen before being perfused with ice-cold PBS then 4% PFA/PBS.

### Transient MCAO

Using aseptic techniques, anesthetized mice (2-3% isoflurane/oxygen) were maintained on a warming pad and an intraluminal filament was inserted into the left MCA via the carotid artery, similar to previously published procedures^52^. A laser Doppler probe (Perimed) monitored blood flow. After 30 minutes of occlusion, the filament was removed. Sterile lactated ringers solution (1 mL SC daily-McKesson) and buprenorphine (0.05 mg/kg IP twice daily) was administered for 48h post-surgery.

### Modified Neurological Severity Score

Neurological impairment was measured by a blinded female researcher using the modified neurological severity score (mNSS), similar to published procedures^53^. Mice lacking deficits score a 0, while the most severe deficits score a 14.

### Immunohistochemistry

After incubating in 4% PFA/PBS ON at 4°C post-perfusion, brains were placed in 30% sucrose/PBS for at least 3 days. Tissue was immersed in OCT (Optimal Cryo-Temp, Tissue Plus – Fisher HealthCare) and frozen using isobutane surrounded by liquid nitrogen. Coronal slices (30 μm) were mounted to gelatin coated slides, dried overnight, and stored at −80°C.

Slides were dried overnight, and then rehydrated in PBS (5 min) prior to permeabilization in triton-X (0.2% in PBS, 10 min). Slides were washed with PBS (2X5min). Autofluorescence was blocked with Sudan Black (0.1% in 70% EtOH, 5 min) and washed with PBS until clear. Non-specific protein binding was blocked with BSA (5% in PBS, 45 min). Primary antibodies were diluted in blocking solution, and tissues were incubated overnight at 4°C. Slides were washed with PBS (4X15 min) before incubation in species-specific secondary antibodies in blocking solution (1 hour RT). Slides were washed in PBS (2X5 min), stained with DAPI (1:10,000 for 5 min, prepared in PBS and stirred for at least 30 minutes), washed with PBS (1×5 min) and then distilled water (1X1 min). Slides were dried and mounted to coverslips with Aqua-Poly/Mount (18606-20).

Antibodies used were rat anti-CXCR4 (R&D MAB21651 1:25); chicken anti-GFAP (Sigma AB5541 1:500); donkey anti-chicken Alexa 488 (Jackson 703-545-155 1:200); donkey anti-rat Alexa 488 (Jackson 711-545-152 1:200).

### Image analysis

Images were obtained using a Zeiss LSM710 laser scanning confocal microscope or Zeiss Axio Observer.D1. Image analysis was performed by a researcher blinded to genotype or experimental manipulation using ImageJ containing the Fiji plug-ins suite.

To measure CXCR4 expression intensity, tdTomato images were used to create region of interest (ROI) selections to limit analysis to Cre+ NSCs: 8-bit images were subject to contrast enhancement to visualize cells, despeckle was used (2X) to remove single pixels, and then was thresholded to include only red signal. The “analyze particles” function created cell masks (>20 pixels). Masks were dilated (2X) and then used to create a selection to add to the ROI manager. ROIs from the tdTomato channel were used to select the same area in the 8-bit CXCR4-positive channel, and the “measure” function measured gray values. For each mouse, z-stacks from at least 3 separate images were measured, and the results are pooled from n=5-6 individual mice.

For *in vivo* migration, the distance between the SVZ and the furthest migrating neuroblast in the coronal plane was measured from the edge of the dorsolateral corner of the SVZ along the center of the migratory track cells to the furthest tdTomato positive cell. Results are pooled from of 3-5 slices/animal of n=13 mice/group.

### RNA extraction and qPCR

Similar to previously published protocols^54^, the SVZ’s (n=4) were micro-dissected and suspended in ice cold hibernation buffer [30mM potassium chloride (CAT# BP366, Fisher), 5mM sodium hydroxide (CAT# S320, Fisher Scientific), 5mM sodium phosphate monobasic monohydrate (CAT# BP330, Fisher Scientific), 0.5mM magnesium chloride hexahydrate (CAT# M2670, Sigma), 20mM sodium pyruvate (CAT# P5280, Sigma), 5.5mM d+ dextrose anhydrous (CAT# D16, Fisher) and 200mM d-sorbitol (CAT# S1876, Sigma)] and dissociated with 10 units of papain and 10μL of DNase in DMEM (37°C for 30 minutes). The cells were triturated, washed, and pelleted then resuspended in FACS Buffer (30% Glucose, 10% BSA in HBSS) and sorted into RNA Later (Thermo-Fisher) using BD FACSAria to select only tdTomato+ cells. RNA was harvested according to manufacturer’s instructions using the RNeasy Micro kit (Qiagen). First-strand synthesis was achieved using the Applied Biosystem High-Capacity cDNA reverse transcription kit according to manufacturer’s instructions. QPCR was run in triplicate reactions using iTaq SYBR Green Supermix (BioRad) according to manufacturer’s instructions on Bio-Rad CFX96 Touch Real-Time PCR detection system. Relative levels of mRNA were determined after normalizing loading via the delta-delta Ct method.

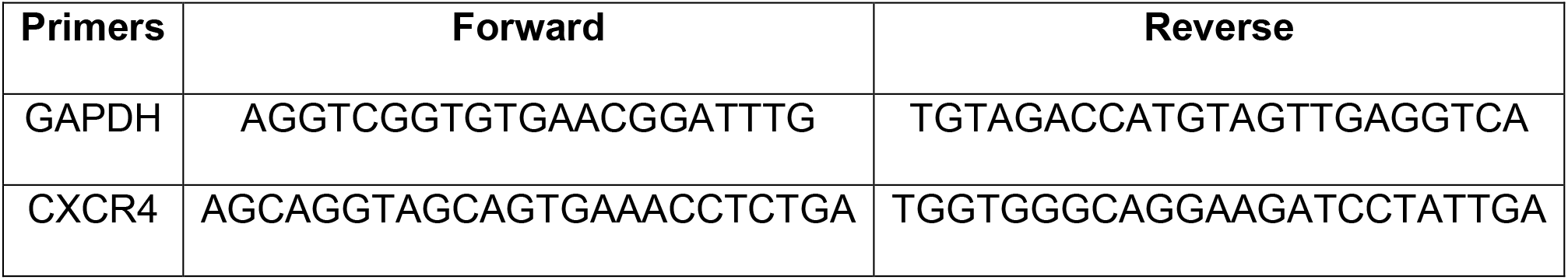

### Chemotaxis Assay

Acutely dissociated SVZ neural stem cells were collected and processed for cell culture similar to previously published protocols^55^. A total of 21,600 cells were plated onto the upper wells (n=6/condition) of 96-well poly-l-ornithine (PLO) coated chemotaxis chambers (10 μm pore size, Neuro Probe Inc) and maintained in DMEM containing l-glutamine (1 mM), sodium pyruvate (1 mM), B-27 (1X from Gibco), N2 (1X from Gibco). The bottom chamber of the wells was loaded with either serum free media or serum free media with SDF1 (500nM). Cells were incubated overnight (37°C, 5% CO2), and then the migrated tdTomato positive cells were counted. Recombination rate was determined by counting the ratio of tdTomato positive/DAPI positive cells from NSCs cultured in-tandem on PLO coated Terasaki plates. Migrated cells were normalized using the recombination rate. Results are pooled from triplicate independent experiments and were analyzed by a researcher blinded to experimental genotype.

### Statistical Analyses

All experiments and analyses were performed by a researcher who was blinded to the genotype/experimental manipulation. Relevant numbers of mice used for each experiment are noted as appropriate. Males and females were used in all conditions. For image analysis, at least 3-4 sections spaced every 150 μm were analyzed and pooled per mouse. For *in vitro* analysis, experiments were performed in triplicate independent experiments with n=3+ wells/condition. Students t-test was used for two-group comparisons and ANOVA with Tukey’s HSD was used to compare more than two groups with α set to 0.05. GraphPad Prism software was used for all statistical analyses. All results show the mean with standard error.

## Acknowledgements

This project was funded by the William and Ella Owens Medical Research Foundation. KD was supported by a training grant supported by the National Center for Advancing Translational Sciences, National Institutes of Health, through Grant TL1 TR002647 and through the American Heart Association Award 916018 co-funded by the Voelcker Fund. Some images were generated in the Core Optical Imaging Facility which is supported by UT Health San Antonio and NIH-NCI P30 CA54174. Data was also generated in the Flow Cytometry Shared Resource Facility, which is supported by UT Health, NIH-NCI P30 CA054174-20 (CTRC at UT Health) and UL1 TR001120 (CTSA grant).

## Author Contributions

KD, SM, JV, SH, EK, and NLS performed experiments, analyzed data, and edited the manuscript. PR and SS performed surgical manipulations and assisted with experiments and manuscript editing.

## Declaration of Interests

None

## Graphical Abstract

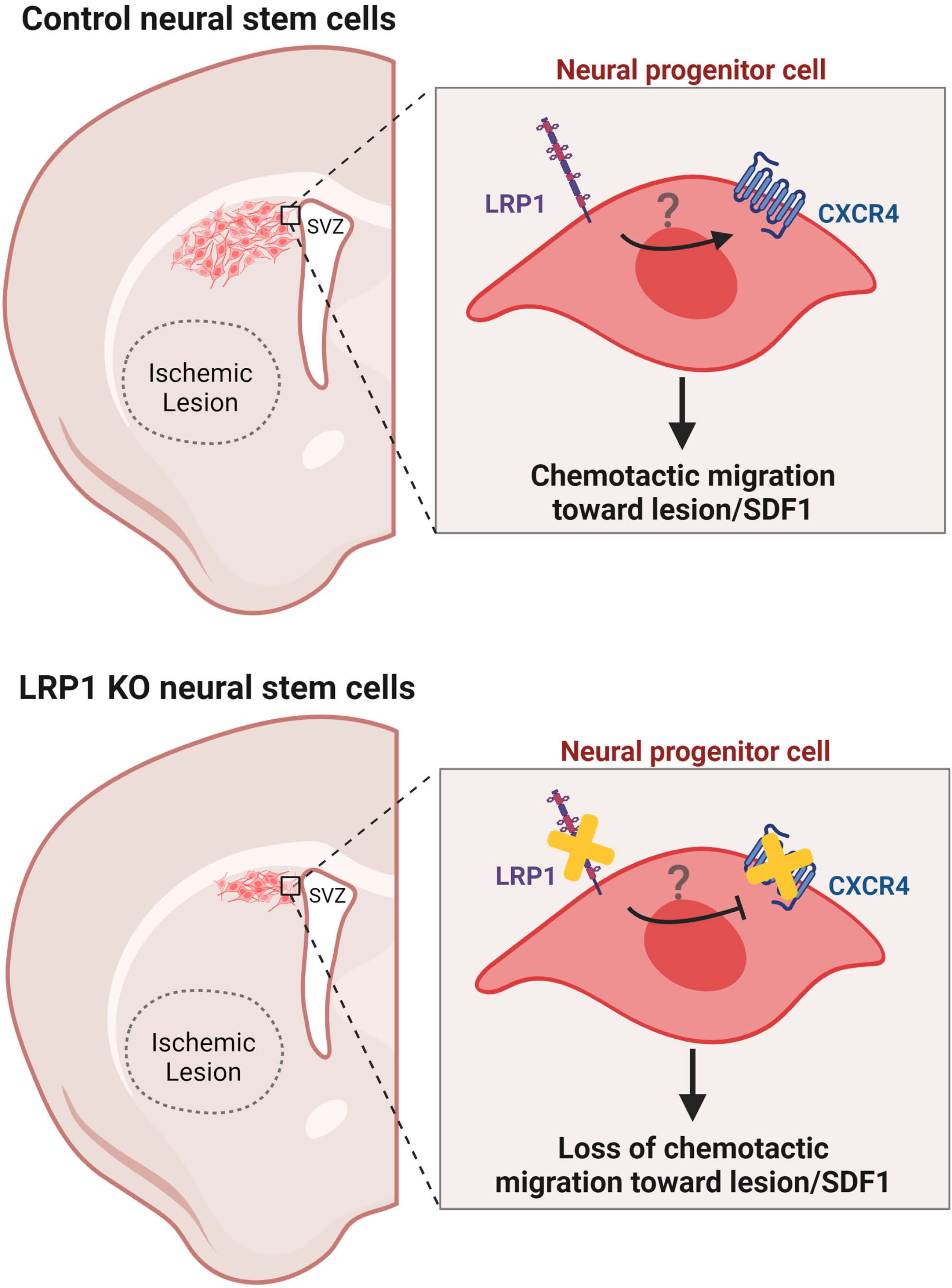

## Notes

### Competing Interest Statement

The authors have declared no competing interest.

